# Identification of clade-defining single nucleotide polymorphisms for improved rabies virus surveillance

**DOI:** 10.1101/2023.08.25.553658

**Authors:** Ankeet Kumar, Sheetal Tushir, Yashas Devasurmutt, Sujith S Nath, Utpal Tatu

## Abstract

**Background:** Rabies is an ancient disease that remains endemic in many countries. It causes many human deaths annually, predominantly in resource-poor countries. Over evolutionary timelines, several rabies virus (RABV) genotypes have stabilised, forming distinct clades. Extensive studies have been conducted on the origin, occurrence and spread of RABV clades. Single nucleotide polymorphisms (SNPs) distribution across the RABV genome and its clades remains largely unknown, highlighting the need for comprehensive whole-genome analyses.

**Methods:** We accessed whole genome sequences for RABV from public databases and identified SNPs across the whole genome sequences. Then, we annotated these SNPs using an R script, and these SNPs were categorised into different categories; universal, clade-specific, and clade-defining, based on the frequency of occurrence.

**Results:** In this study, we present the SNPs occurring in the RABV based on whole genome sequences belonging to 8 clades isolated from 7 different host species likely to harbour dog-related rabies. We classified mutations into several classes based on their location within the genome and assessed the effect of SNP mutations on the viral glycoprotein.

**Conclusions:** The clade-defining mutations have implications for targeted surveillance and classification of clades. Additionally, we investigated the effects of these mutations on the Glycoprotein of the virus. Our findings contribute to expanding knowledge about RABV clade diversity and evolution, which has significant implications for effectively tracking and combatting RABV transmission.

## Introduction

Rabies virus (RABV), along with some other species of the genus *Lyssavirus*, causes a disease called rabies. It is a deadly zoonotic viral encephalitis that is fatal in almost 100% of cases and significantly threatens human and animal health [1]. The *Lyssavirus* genus comprises 17 species, including the RABV, type species of the genus and other rabies-related viruses [2]. These species are classified into phylogroups based on genetic composition and antigenic differences [3]. Most Lyssavirus species have a limited impact on human health due to their narrow host range. RABV displays broad host tropism and is responsible for approximately 59,000 human deaths annually across the globe [4]. Almost 99% of human deaths caused by RABV globally are attributed to dog bites [1]. In India, dogs are the only known reservoir of RABV, and India bears the highest burden of rabies-related human deaths, accounting for around 20,000 fatalities annually [5], followed by China [6].

The RABV genome is approximately 12 kilobases long. The genome is non-segmented, single-stranded negative-sense RNA enclosed within an enveloped, bullet-shaped virion [7]. The genome codes for 5 proteins are arranged in the following order: Nucleoprotein (N), Phosphoprotein (P), Matrix (M), Glycoprotein (G), and Large (L) protein genes. The L gene is the largest, comprising approximately 53% of the total nucleotide content. It codes for a crucial, multifunctional protein called RNA-dependent RNA polymerase (RdRp) [8]. Like most RNA viruses, RABV RdRp lacks proofreading activity [9], resulting in variations, ultimately leading to the emergence of new variants and the formation of quasi-species during infection [10]. P protein codes for various isoforms; its full-length P protein interacts with the RdRp protein via its N-terminal region, while its shorter isoforms (P2-P5) with truncated N-termini likely serve alternative functions [11]. The N protein forms the core of the nucleocapsid, encapsulating the viral RNA in a helical structure [12]. The M protein interacts with the N protein and G protein facilitating viral envelopment [13]. The G protein is a trimeric and is essential for viral entry [14]. The G protein is known to interact with multiple host receptors [15]. It is also a target for neutralising antibodies and a therapeutic target in the virus [16].

Previously, studies primarily relied on sub-genomic sequences[17,18], which has constrained our ability to understand whole-genome-level attributes comprehensively. Additionally, to our knowledge, there is no systematic study tabulating the global occurrence of single nucleotide polymorphisms (SNPs) in the RABV and comparing SNPs across the clades; a notable gap exists in the characterisation of clades. In this study, we present a comprehensive analysis of SNPs based on whole genome sequences of 8 clades of RABV isolated from 7 host species (Dog, Cat, Cattle, Horse, Sheep, Goat and Human). The SNPs result in various types of mutation at the protein level. The percentage of these mutations differs across the clades. We report some mutations occurring with high frequency (clade-specific, universal and clade-defining) and further analyse their impact on the G protein.

## Methods

### Collection and Curation of the WGS Data

To identify SNPs in the WGSs of the virus, we accessed data from 2 databases-Bacterial and Viral Bioinformatics Resource Center (BV-BRC) [19] and RABV-GLUE (RABV-GLUE) [20] (databases accessed on 1/11/2023). The databases contained varying numbers of WGSs for the representative hosts. Comprehensive datasets were generated through manual curation, which involved merging non-overlapping sequences from two databases and removing redundancies. Annotations were further verified against NCBI databases (Nucleotide and Bio-sample). The pipeline of manual curation is presented in S1 Fig.

This study focuses on the dog-variant of RABV; therefore, we downloaded sequences isolated from the Dog host and 6 other hosts likely to harbour the dog variant of RABV. The other hosts were cat with 43 sequences, cattle with 106 sequences, horse with 8 sequences, sheep with 12 sequences, goat with 23 sequences and human with 47 sequences.

Sequence metadata files were downloaded from both databases. Metadata lacking clade information was supplemented using RABV-GLUE, a web-based tool from the University of Glasgow [21]. Only relevant information was retained in the refined metadata. Clade distribution across different hosts and the corresponding genome count is presented in Table 1.

**Table 1.** Dataset of hosts and their clades with the associated count.

| Host Number | Host Name | Clade Number | Major Clade | Number of genome sequences |
| --- | --- | --- | --- | --- |
| 1 | Dog (*Canis lupus familiaris*) | 1 | Cosmopolitan | 383 |
|  |  | 2 | Asian | 65 |
|  |  | 3 | Arctic | 50 |
|  |  | 4 | Africa-2 | 37 |
|  |  | 5 | Bats | 3 |
|  |  | 6 | Indian subcontinent (Indian-Sub) | 1 |
| 2 | Cat (*Felis catus*) | 1 | RAC-SK (Raccoon-Skunk) | 18 |
|  |  | 2 | Cosmopolitan | 13 |
|  |  | 3 | Africa-3 | 3 |
|  |  | 4 | Arctic | 3 |
|  |  | 5 | Asian | 3 |
|  |  | 6 | Africa-2 | 2 |
|  |  | 7 | Bats | 1 |
| 3 | Cattle (*Bos taurus*) | 1 | Cosmopolitan | 70 |
|  |  | 2 | Arctic | 12 |
|  |  | 3 | RAC-SK | 11 |
|  |  | 4 | Bats | 7 |
|  |  | 5 | Asian | 5 |
|  |  | 6 | Indian-Sub | 1 |
| 4 | Goat (*Capra hircus*) | 1 | Cosmopolitan | 22 |
|  |  | 2 | Arctic | 1 |
| 5 | Horse (*Equus caballus*) | 1 | RAC-SK | 3 |
|  |  | 2 | Cosmopolitan | 2 |
|  |  | 3 | Bats | 1 |
|  |  | 4 | Asian | 1 |
|  |  | 5 | Arctic | 1 |
| 6 | Human (*Homo sapiens*) | 1 | Cosmopolitan | 14 |
|  |  | 2 | Asian | 13 |
|  |  | 3 | Arctic | 8 |
|  |  | 4 | Africa-2 | 6 |
|  |  | 5 | Indian-Sub | 4 |
|  |  | 6 | Bats | 2 |
| 7 | Sheep (*Ovis aries*) | 1 | Asian | 7 |
|  |  | 2 | Cosmopolitan | 4 |
|  |  | 3 | Arctic | 1 |

The final metadata file for 778 genome sequences utilised in the study is present in the S1 File.

### Identification of Single Nucleotide Polymorphisms

The identification of SNPs involved several steps. The multi-fasta sequence files for hosts were stored in different directories. These fasta files were subjected to SNP analysis using the NUCMER v3.1 [22]. This tool generates pairwise alignment of each sequence against the reference sequence of RABV (NC_001542.1) and produces a table of SNPs. The process was repeated for each host. This SNP table contained information regarding the variant position and nucleotide change compared to the reference sequence. The table generated from Nucmer was used for further analysis using R (v4.3.1).

### Classification of Single Nucleotide Polymorphisms using R programming

The original R script for SNP analysis [23] was modified (S2 File) for improved accuracy by changing the “round” function to “ceiling”. This was necessary because the “round” function could lead to misinterpretation of changes occurring at the first nucleotide position of a codon. This script utilises the SNP table generated using Nucmer in the previous step. The script utilises a General Feature Format 3 (GFF3) file containing information about the start and end positions of the genes. The GFF3 file for the RABV reference genome is available as S3 File. This GFF3 was manually curated to keep the information only for coding regions. The script utilises Seqinr (v4.2.30) [24] and Biostrings (v2.68.1) [25] tools to load a reference fasta file and translate the nucleotide changes into amino acids using the reference genome backbone. The translated amino acid changes are labelled as SNP (non-synonymous change) mutation, SNP-silent (synonymous change) mutation, deletion-frameshift, insertion frameshift and SNP-stop. The region falling outside the genes is labelled as extragenic. The generated data frame containing amino acid changes includes duplicate rows for an amino acid where more than one nucleotide change is responsible. To eliminate these duplications, an R script (S4 File) was written to merge rows with multiple nucleotide changes corresponding to the same amino acid residue.

### Categorisation of High-Frequency Mutations

Most observed mutations were occurring with low frequencies. However, our analysis focused on high-frequency mutations because of their potential significance in the biology of the virus. Initially, we pinpointed mutations prevalent within individual hosts. The mutations appearing in over 90% of sequences within a given host were denoted as “universal mutations”. Subsequently, given the presence of multiple clades within each host, our attention turned to “clade-specific mutations”. These mutations represented unique and non-overlapping mutations localised within distinct clades coexisting within the same host. To ensure the specificity of clade-specific mutations, universal mutations were excluded from the dataset during the process. Finally, as similar clades were found in more than one host, we discerned “clade-defining mutations” by identifying common clade-specific mutations across different hosts for a clade. These mutations, characterised by their high frequency and consistent presence, served as distinctive markers reliably delineating specific clades.

Due to the reliance on percentage values for classification, a minimum threshold of four sequences in a host and the presence of at least two clades for comparison were established as criteria for inclusion in the high-frequency mutation analysis. Consequently, horse and goat data were excluded from this analysis as they failed to meet the criteria (S2 Fig.).

### Assessing the Impact of Mutations on the Glycoprotein

To elucidate the mutational effects on the structure of RABV glycoprotein, homology modelling and protein threading techniques were implemented. The RABV glycoprotein backbone was modelled using the post-fusion Mokola virus structure (PDB ID: 6TMR), while the stalk region utilised the H5N1 influenza virus hemagglutinin (PDB ID: 4UJM) as a template, both in SWISS-MODEL [26]. DynaMut [27], which provides computational predictions on the impact of mutations on proteins, was employed to predict the impact of some of these mutations on protein.

### Data Visualisation and Bioinformatics Tools

We utilised R (v4.3.1) for data analysis and visualisation. A map was created to understand the global distribution of the RABV clades found in dog hosts (Fig. 1A) using the Maps v3.4.1 tool from the R library. Genetic diversity at the nucleotide level was calculated using DnaSP6 program [28]. The calculation of average mutations for the 5 proteins of RABV clades is done using base R functions and dplyr package v1.1.2. The count of mutation is normalised to protein length and averaged by sample number for that clade. The library drawProteins v1.20.0 [29] was used to draw domains for different proteins in Fig. 5A. The structure was visualised in the PyMol and ggplot2 v3.4.2 [30] was utilised to make figures.

**Fig 1.**
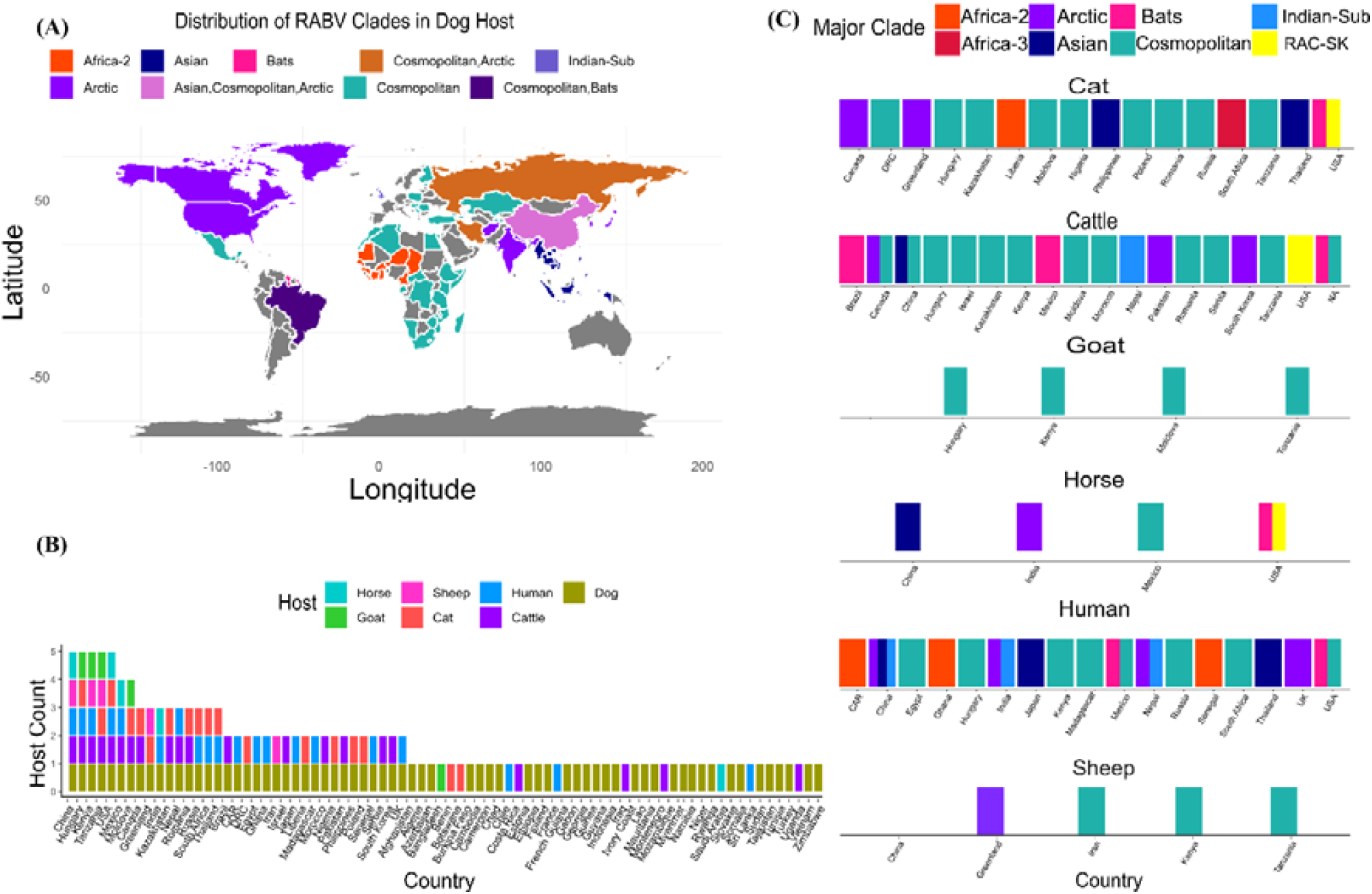
Geographic distribution of different clades of RABV based on whole genome sequences (N=778). **A**. The world map shows the prevalence of RABV clades found in the dog host; colours highlight different clades. **B**. Distribution of hosts across 78 countries. The coloured bars represent different hosts. **C**. Represents countries with more than one host or a single host with more than one clade. Major clades are represented in different colours

## Results

### Global Distribution and Host Association of Rabies Virus Clades

Rabies virus shows global presence and, over time, has diversified into distinct genotypes forming monophyletic clades [31]. Fig. 1A displays the spatial distribution of the 6 RABV clades isolated from the dog hosts. The cosmopolitan clade demonstrates the broadest spatial spread. Most countries predominantly harbour a single clade; China exhibits 3 clades, and Russia, Iran, and Brazil show 2 clades each. The Asian clade ranks second in numbers for dog hosts in our dataset; the clade was reported from China and neighbouring Southeast Asian countries. The Arctic clade spans three continents (Europe, Asia, and the USA). However, dog-mediated rabies is not endemic to America and Europe [32]. The Africa-2 clade is restricted to western Africa. The Indian-Sub clade is reported from the UK, and the Bats clade in dogs is found in South American countries.

Fig. 1B shows the occurrence of hosts in 78 countries. None of the countries in the dataset showed the sequencing data for all 7 hosts simultaneously. The maximum number of hosts observed in any single country was 5-China, Hungary, Kenya, Tanzania, and the USA demonstrated RABV sequences from 5 hosts, followed by Mexico and Moldova, which represented 4 hosts each. Dog hosts were found to be the most common host (observed in 67 countries), followed by cattle (in 22 countries), humans (in 21 countries), and cats (in 18 countries). Conversely, goats, horses, and sheep were the least common, each represented in only 5 countries. Forty-three countries had data only from 1 host; out of these 43 countries, around 72% showed sequences only from dogs.

To understand the co-occurrence of hosts and clades in the dataset, we looked for countries with either more than 1 host or more than 1 clade; most of the countries passed the criteria, while Indonesia and Bangladesh had only one host harbouring a single clade, so they were excluded. As shown in Fig. 1C, the Cosmopolitan clade is the most common clade across all 7 hosts, followed by the Arctic. Bats clade is reported from the USA, Mexico, and Brazil, and RAC-SK is only reported from the USA.

### Characterisation of Mutation Classes and Nucleotide Position Biasness in RABV Genome Across Diverse Hosts

RNA viruses show a high mutation rate. To understand the differences in RABV clades at the molecular level, we identified nucleotide variations and categorised them into six classes: SNP-silent, SNP, SNP-stop, extragenic, insertion-frameshift and deletion-frameshift mutation using an R script. A detailed breakdown of mutation counts categorised by functional class for each RABV clade in the five proteins and the extragenic region, with respect to different host species, can be found in the S1 Table. Fig. 2A presents the percentage of various classes of mutations based on 539 dog RABV sequences belonging to 6 clades (Africa-2, Arctic, Asian, Bats, Cosmopolitan, and Indian-Sub). The synonymous mutation or SNP-silent is the most prevalent (70-82%) class of mutations across the clades. The subsequent mutation classes are SNP mutation and extragenic mutation class, which account for most of the remaining variations (approximately 30%). Less than 1% of mutations fall into insertion-frameshift and deletion-frameshift classes. The percentage of non-synonymous mutation in different clades ranges between 11-15%. The extragenic class ranged from 6-15% across the clades. The cosmopolitan clade displayed the highest percentage of extragenic and SNP mutations in all the hosts. The reason for displaying higher SNP mutations in cosmopolitan could be because multiple sub-clades have become associated with geographical locations.

**Fig 2.**
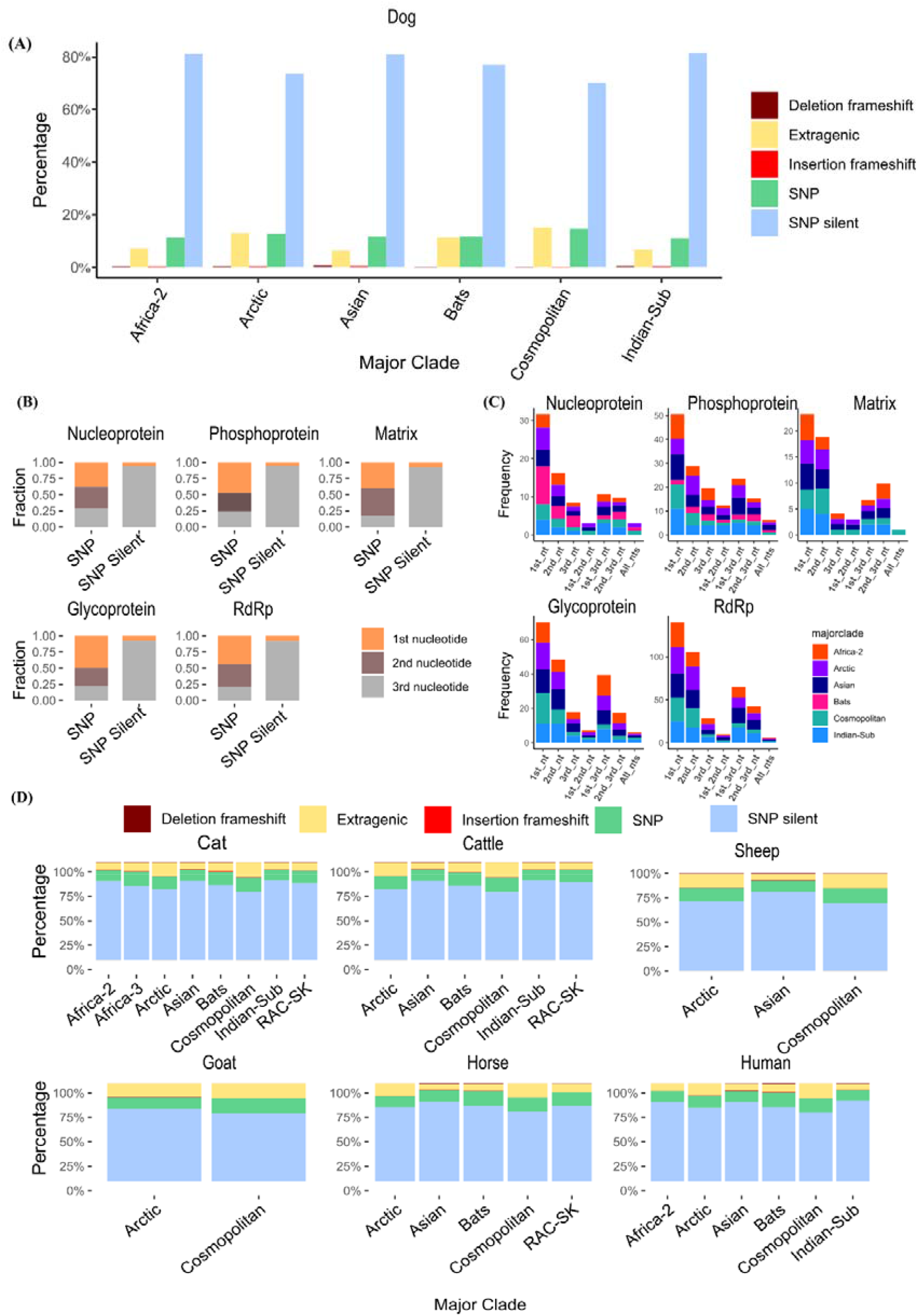
Analysis of mutation classes and underlying nucleotide positions involved in RABV. **A**. The percentage of different mutation classes occurring in the RABV clades of dog host. **B**. The contribution of nucleotide position within a codon for the SNP and SNP silent mutation class in RABV genes. The y-axis contains the fraction contributed by three nucleotides of a codon. **C**. The fraction of different (individual and co-occurring) nucleotide positions to SNP mutations in the RABV genes. **D**. The percentage of mutation classes observed in different clades of the other six hosts.

Additionally, we analysed the contribution of the nucleotide positions within a codon on SNP and SNP silent mutations. Fig. 2B shows that SNP silent mutations predominantly result from III nucleotide change within a codon. Only a minor fraction of SNP silent mutations occur due to changes in I nucleotide and even less due to II. Conversely, all three nucleotide changes contribute to SNP mutations significantly. Both I and II nucleotide changes contribute almost equally to SNP mutations in RABV proteins except for G and P, which exhibited lower contributions from II nucleotide change. While SNPs primarily arise from individual nucleotide changes at the I and II positions of codons, a considerable portion also results from simultaneous substitutions within a single codon. The co-occurrence of SNPs within a codon depicted contribution disparity within clades and proteins, as shown in Fig. 2C.

Fig. 2D represents the occurrence of various classes of mutations in clades of other hosts present in the dataset (Cat, Cattle, Sheep, Goat, Horse and Human). Like dogs, the variations observed in other host species also showed SNP silent as the predominant mutation, followed by SNP and extragenic mutation. The Cosmopolitan clade demonstrated the lowest percentage of SNP-silent mutations throughout all hosts compared to other clades. These results highlight differences in the selection pressures across the RABV clades and hosts. Data on the mutations found for different hosts is presented in the S5 File.

### Phosphoprotein Exhibits Highest Mutation and C>T is Major Transition Across Diverse Clades of RABV

Previous studies have reported that the Phosphoprotein shows the highest mutation rate [33]. We investigated the mutation patterns across diverse RABV clades present in 7 hosts. Fig. 3A depicts mutation trends across 5 proteins of RABV clades isolated from dogs. In our study, most clades showed Phosphoprotein as the highest mutating protein among the 5 RABV proteins in the sequences isolated from dogs. The Glycoprotein and RdRp exhibited similar average mutations despite the huge difference in protein length. Notably, the Bats clade exhibited a distinct pattern of variation, with the highest number of mutations observed in the Nucleoprotein, followed by fewer mutations in the Phosphoprotein, and no mutations detected in the Matrix, Glycoprotein, and RdRp proteins.

**Fig 3.**
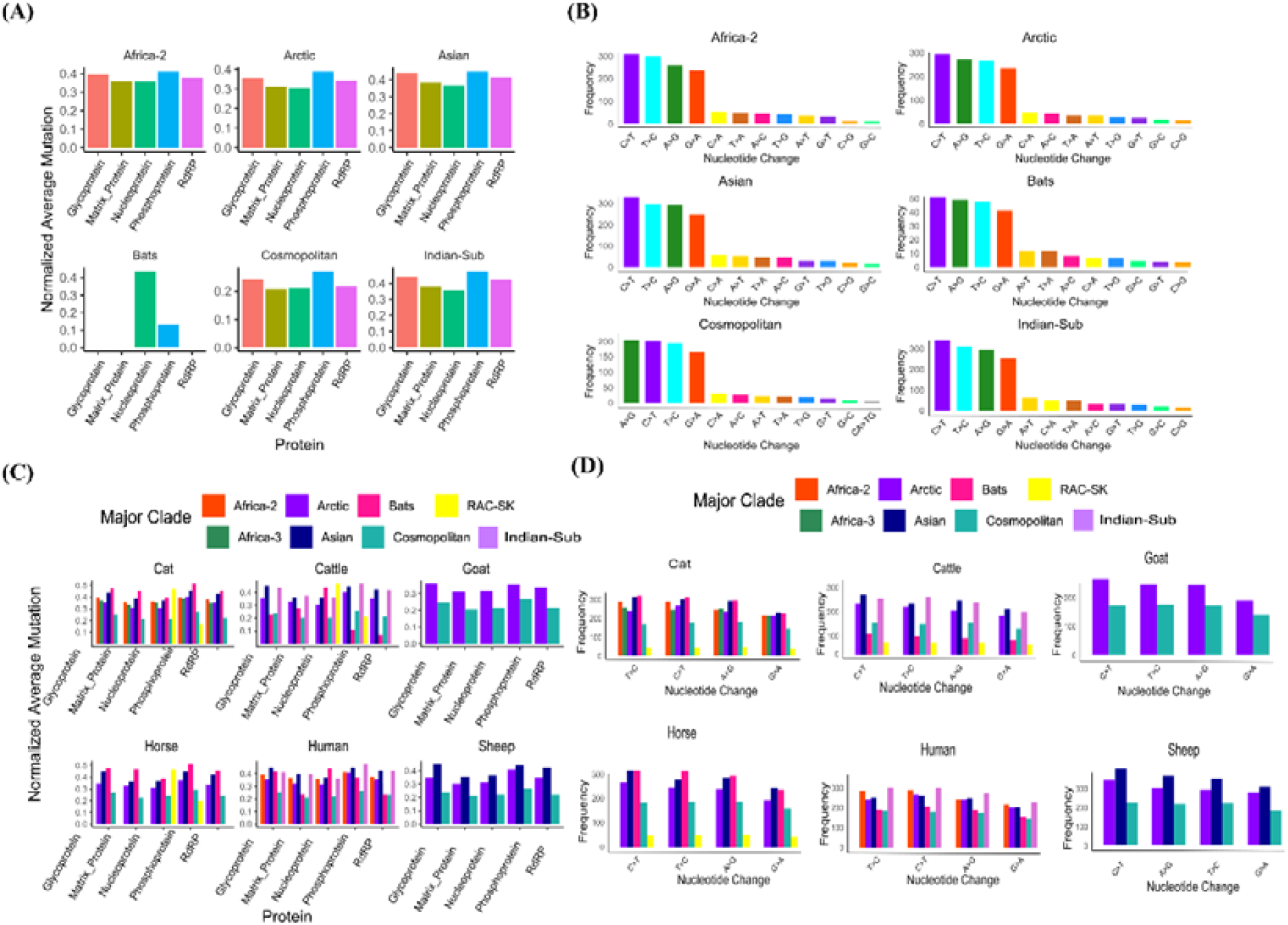
Mutation counts across five proteins of RABV and underlying nucleotide transitions and transversions. **A**. Mutation count normalised over protein length in the RABV proteins. The x-axis presents the proteins of RABV, and the y-axis represents the mutation count normalised to the protein length. **B**. The top 12 most common nucleotide transition and transversion in different RABV clades (dog host). The coloured bars represent nucleotide change. **C**. Mutations observed for RABV proteins in other hosts. **D**. Representation of the four most common nucleotide transitions in the clades of other hosts.

The nucleotide transitions and transversion across different clades found in the dog host are presented in Fig. 3B. The top 4 nucleotide transitions (C>T, A>G, T>C, G>A) were consistent throughout the clades. Cytosine to Thymine (C>T) was found to be the predominant transition in 5 clades, while A> G was found to occur with the highest frequency in the Cosmopolitan clade. Transversions were present, but the frequency was lower than the transitions. The trend of these changes varied across the clades in dogs.

Fig. 3C delineates mutations across 5 proteins within diverse RABV clades found in 6 hosts. While the overall mutation trend (P>G∼L>M>N) is the same as that found for the dog host, distinct patterns emerged in the hosts harbouring Bats and RAC-SK clades. For instance, the Bats clade showcased heightened N protein mutations in the cattle and human hosts, whereas P protein displayed elevated mutations in horse and cat hosts. In the cattle host, the mutations were significantly less in Phosphoprotein and RdRp for the Bats clade. The RAC-SK clade consistently exhibited more mutations in the N protein across all the hosts. Interestingly, no mutations were observed in the Glycoprotein and Matrix proteins across any host for the RAC-SK clade, while mutations were seen for RdRp in the cattle host.

The 4 most common nucleotide transitions found in dog RABV were checked in other hosts as well (Fig. 3D). C>T transition was dominant in sheep, goat, cattle, and horse hosts. However, the T>C transition prevailed in human and cat hosts. In the sheep, host A>G was the second most common nucleotide transition, while in all other hosts, it was third. Collectively, these results highlight the different evolutionary pressures across RABV clades in different hosts.

### Mapping the Distribution of Rabies Virus Mutations Across Clades and Identifying Universal Mutations

Similar to other RNA viruses, RABV lacks a proofreading mechanism which drives its ability to accumulate mutations quickly [34]. Fig. 4A sheds light on the distribution of mutations identified across the four RABV clades circulating in dog hosts. The analysis revealed that the Cosmopolitan clade harboured the most mutations, followed by Asian, Arctic, and Africa-2 clades in dogs. This trend also remains consistent for unique and non-overlapping mutations within each clade. These findings suggest that the Cosmopolitan clade may have undergone greater evolutionary pressure or higher adaptability, facilitating its survival in diverse environments. Moreover, the Cosmopolitan clade exhibits the highest degree of overlap with the Asian clade, sharing 489 mutually, followed by the Arctic clade. Notably, 1539 mutations are common across all four clades of dogs and most of these mutations primarily occur at low frequencies (data not shown). Fig. 4B depicts the distribution of universal mutations found across 5 hosts. The sheep host displays the highest unique universal mutations, followed by dog, cat, human and cattle hosts. Sheep possessed 144 unique universal mutations, while 78 were common in dogs and sheep. Human hosts shared 2 universal mutations with dogs, and 1 mutation was exclusive to human hosts. Cattle and cats shared no mutation exclusively with the dogs. A total of 9 mutations were found to be conserved across all the hosts, potentially essential for viral functions that remain unaltered across diverse hosts.

**Fig 4.**
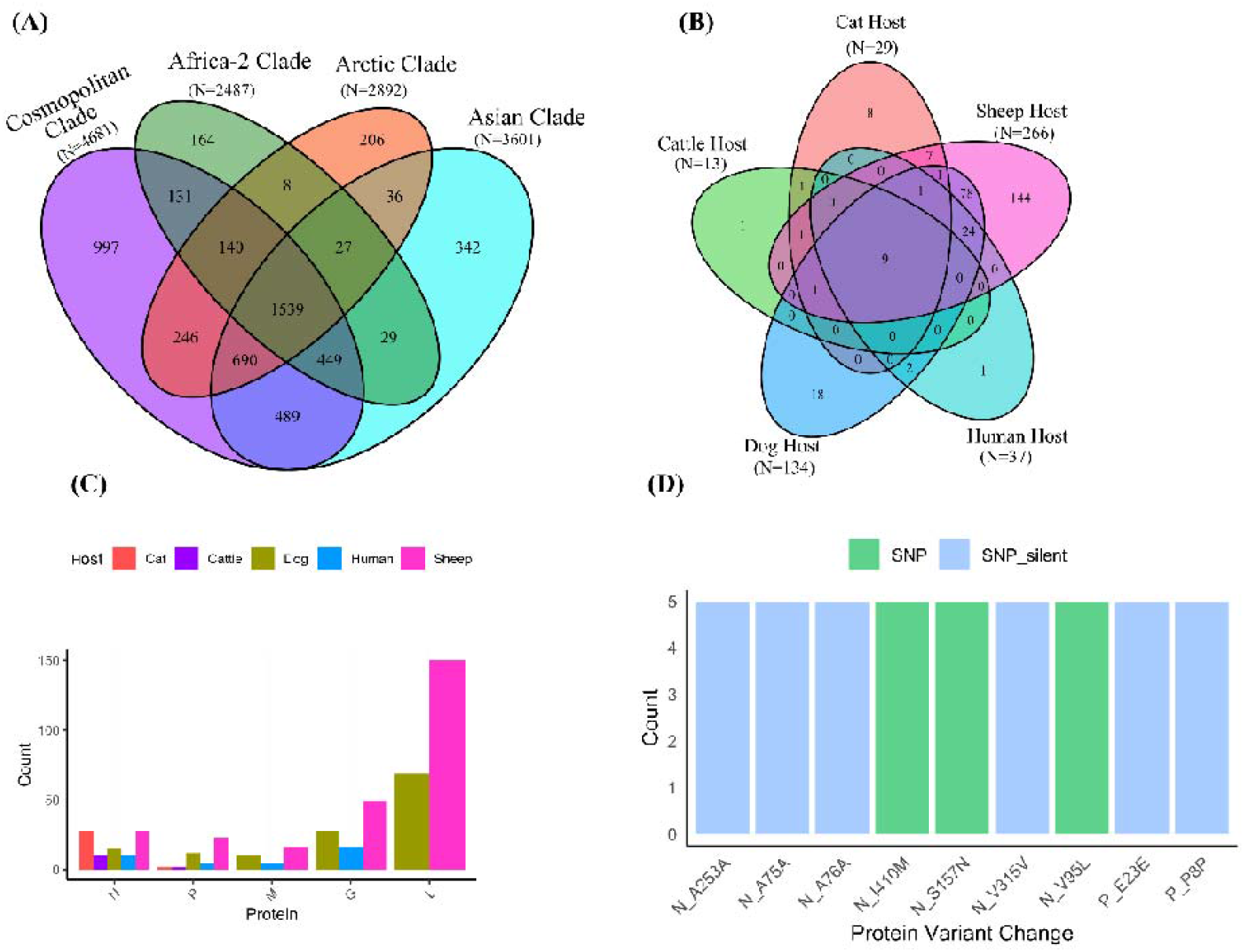
The distribution of mutations in dog RABV and universal mutations in other hosts. **A**. Venn diagram shows the distribution of SNP and SNP silent mutations in RABV clades isolated from dog hosts. The “N” indicates the total number of unique mutations for the clades. **B**. Venn diagram depicts the distribution of universal mutations across five hosts.**C**. Highlights the distribution of universal mutations in the five proteins across different hosts; the colours represent hosts. **D**. Representation of nine common mutations in RABV sequences found across all hosts.

**Fig 5.**
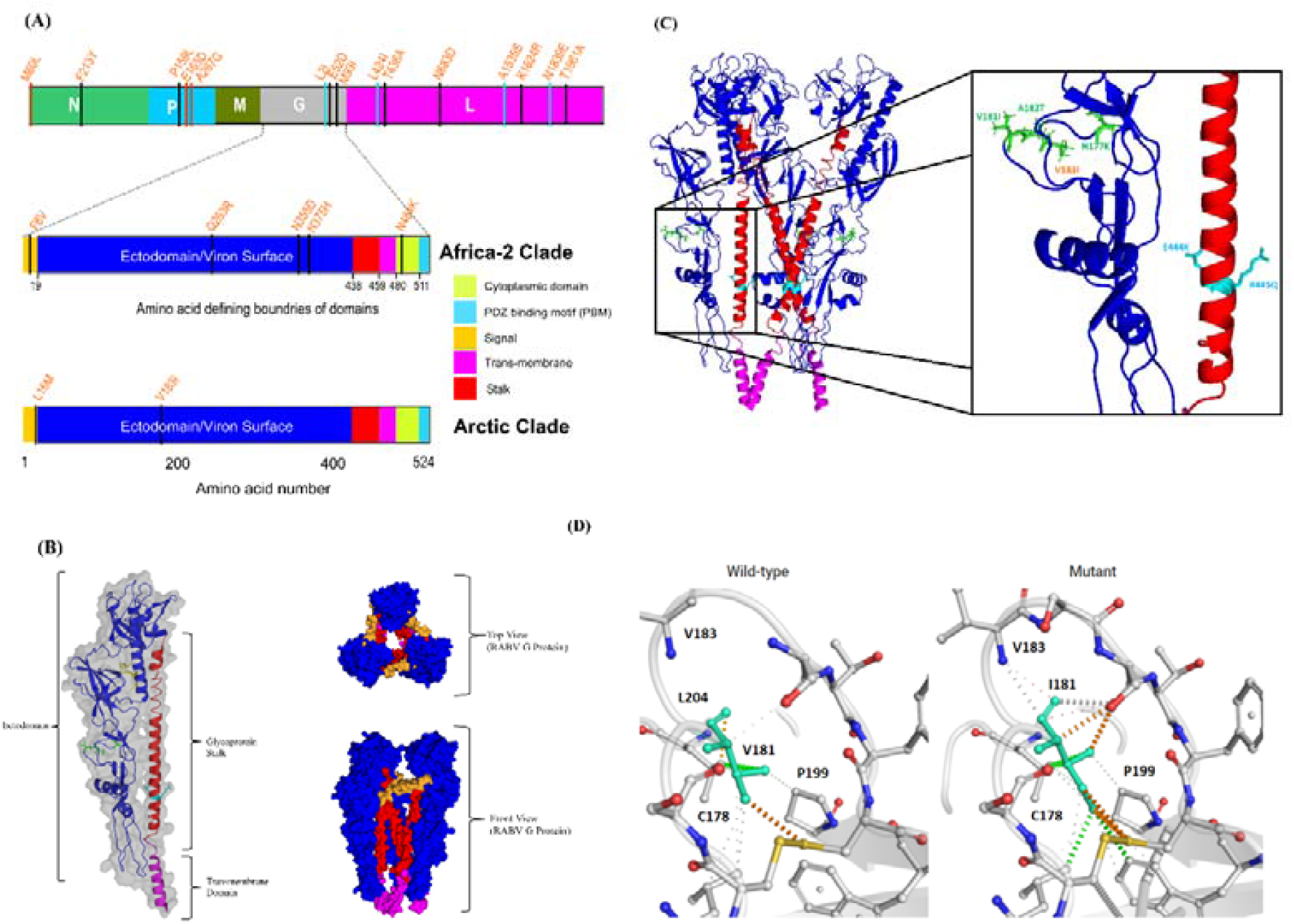
Clade-specific mutations in the glycoprotein. **A**. G protein with different ectodomains, showing clade-specific mutations. **B**. RABV Glycoprotein structure predicted using the SWISS-MODEL, showing the post-fusion form of protein. The transmembrane region is highlighted in pink, the stalk is coloured red, and the ectodomain is represented with blue colour (left). The surface structure of trimeric-RABV Glycoprotein top and front view (right). **C**. The trimeric ribbon representation showing the interaction of three monomers of Glycoprotein (left). Zoomed-in box represents mutations occurring in the AchR receptor binding region. The green and orange represent universal and clade-specific mutations, respectively. Two universal mutations, E444K and R445Q occur in the stalk region of the glycoprotein. **D**. Represents the alteration of the interaction upon V to I change at 181 residue.

The distribution of these universal mutations across proteins is showcased in Fig. 4C. RdRp protein displayed the highest universal mutations, but the mutations were only seen for dog and sheep hosts. This was followed by G and M proteins, which showed mutations for dog, human and sheep hosts. The N and M showed universal mutations for all the hosts. Fig. 4D highlights 9 conserved mutations common to RABV and found across all hosts in high frequency. Only 3 belong to the SNP mutation class, and the rest are SNP silent mutations. This subset of non-synonymous mutations may contribute to subtle functional shifts that enhance viral fitness. A list of unique mutations in hosts with the universal mutations for the respective hosts is provided in the S6 File. This mutation landscape sheds light on the evolutionary dynamics of RABV and suggests that mutation accumulation, particularly in receptor-binding and replication-related proteins, may contribute to the virus’s ability to persist and adapt across different hosts.

### Assessing the Impact of High-Frequency Mutations on RABV Glycoprotein

The RABV glycoprotein is crucial in facilitating virus entry and serves as the primary target for neutralising antibodies, making it a key protein for host adaptation and immune evasion [14]. Fig. 5A highlights the clade-specific SNP mutations present in the 4 proteins. M protein did not display any clade-specific SNP mutation. N protein displayed 1 mutation in Africa and the Arctic each. P displayed 1 and 2 mutations in Arctic and Africa-2, respectively. RdRp protein, essential for viral replication, displayed 5 clade-specific SNP mutations in Africa-2 and 4 in Asian clades. G protein displayed 5 clade-specific mutations in Africa and 2 mutations in the Arctic clade. The distribution of these mutations may clade-specific adaptations and immune evasion strategies. The stability values for mutations in the G protein are available in the S2 Table. A complete list of all the clade-specific mutations observed in dog RABV clades is presented in Table 2. The RdRp protein showed the highest number of clade-specific mutations in dogs, and the mutations in RdRp were found for the Africa-2 and Asian clades. Africa-2 showed clade-specific mutations in all 5 proteins. All the clade-specific mutations seen in Glycoprotein belonged to the SNP class.

**Table 2.** Clade-specific mutations were observed in 3 clades of RABV in dog hosts (bold values highlight SNP mutations).

| Clade | Protein | Mutations |
| --- | --- | --- |
|  | Nucleoprotein | **F213Y** |
|  | Phosphoprotein | **K139K, P159L** |
| Africa-2 | Matrix Protein | S185S |
|  | Glycoprotein | **F8V, Q263R, N355D, N375H, N484K** |
|  | RdRp | **M93I, T436A, N683D, Q745Q, K1624R, T1961A** |
| Arctic | Nucleoprotein | **M60L**, P135P |
|  | Phosphoprotein | **E165D, A267G** |
|  | Glycoprotein | **L16M, V183I** |
| Asian | RdRp | **L2I, L424I, A1535S, N1839E** |

Fig. 5B shows the post-fusion structure of a monomer of Glycoprotein of RABV, with mutations in the stalk and ectodomain region. The surface structure of trimeric Glycoprotein from the top and front view is also presented. Fig. 5C shows the trimeric structure of the Glycoprotein; the zoomed-in portion highlights the 6 mutations occurring in 2 regions. The mutations in blue (ectodomain) occur in the snake-toxin-like region of the Glycoprotein (V181I, A182T, and N177K are universal mutations, while V183I is the only clade-specific mutation occurring in the Arctic clade in this region). 2 universal mutations E444K and R445Q are located in the stalk region of the Glycoprotein, and this region is essential for the interaction of 3 monomers of the Glycoprotein [35]. Fig. 5D highlights the altered interactions between residue 181 and the neighbouring residues, likely impacting protein function. The free energy values suggest it to be one of the highest stabilising mutations, and the interactions are seen to have increased in the mutated version of the residue. This increased stability may enhance viral entry. The clade-specific mutations for other hosts and clade-defining mutations are presented in the S7 File. The accumulation of clade-specific mutations in different proteins across RABV clades may reflect the evolutionary pressures.

## Discussion

Rabies is one of the oldest zoonotic diseases known to mankind. Regardless of the availability of ample WGSs, the characterisation and comparison of SNPs across the clades were lacking. We report SNPs in RABV clades leveraging a dataset comprising 778 whole genome sequences isolated from 7 hosts. Despite the global presence of RABV, the sequencing data available for it is highly biased, severely affected regions lack surveillance regarding the numbers and diversity of the hosts. This leads to the underestimation of infected cases and the types of hosts infected. The re-emergence cases of rabies [36,37] demand robust surveillance mechanisms in dogs as well as other hosts to reach the “Zero by 2030” goal [38].

Most of the mutations identified in this study occurred with a low frequency, yet some low-frequency mutations could still play a crucial role in viral evolution and adaptation. Though these mutations are not extensively described here, their longitudinal tracking is essential to understanding emerging variants. The P protein displayed the highest mutation across most of the clades, likely due to host immune pressure and clade-adaptive strategies. While in some cases for the Bats clade and all the cases for the RAC-SK clade, the mutations were higher in the N protein, emphasising the divergence in mutation pattern due to host-specific factors, although more studies are required to understand these dynamics. SNP silent was found to be the major class of RABV mutations, highlighting the negative selection in RABV and limiting the changes in the protein sequence. Even though the SNP-silent mutations do not change amino acids, the difference at the nucleotide level contributes to changes in codon usage [39,40] and is also known to change protein structure [40], suggesting that these mutations are not entirely inconsequential. The percentage distribution of mutation types varies across the clades. This analysis presents that there are inherent differences in the clades, and their association with the host must be explored to underscore the adaptive mutations of RABV across different clades.

The presence of universal mutations, along with distinct clade-specific mutations, indicates that each clade may require a unique set of mutations for adaptation and survival. These mutations appear to have stabilised within their respective clades over the course of evolution despite the rapid mutation rate of RABV, suggesting their importance in viral biology. Bonnaud et al. (2019) identified adaptive mutations in the N and G proteins of RABV that facilitated the adaptation of dog variant in foxes [41]. They reported that transmission was unidirectional, and several mutations observed in their experiment were seen in the sequences isolated from natural hosts. However, the study only used cosmopolitan strain, and we found that residues described for the adaptation in their study belonged to the universal class in dog hosts (residue 61 in N protein and residue 480 in G protein). This suggests that studying SNPs will help unravel the adaptive mechanisms of the virus and the evolutionary forces responsible for the changes and cross-species transmission.

The clade-specific mutations represent key mutations required for viral adaptation. We compared the clade-specific mutations across the same clade in different hosts and identified clade-defining mutations (S7 File). These clade-defining mutations offer a promising avenue for detecting and categorising clades based on SNPs. We have developed a PCR-based rapid typing tool based on SNPs (data not shown), which will improve the epidemiological tracking of RABV clades by providing an alternative way to determine clades where genome sequencing is inaccessible. A summary table for clade-specific and clade-defining mutations for dog hosts is presented as S4 Table. Similar strategies have been proven effective in tracking viral lineages, notably in SARS-CoV-2 [42]. Our analysis unveils the absence of clade-specific mutations within the cosmopolitan clade. Also, the presence of high-frequency mutations for subclades (S8 File) hints that the diversity at the sub-clade level is higher in the Cosmopolitan clade. The Cosmopolitan showed high extragenic mutations, although extragenic mutations do not directly contribute to the protein sequence but are crucial for protein synthesis and regulation [43].

The genetic diversity calculated for RABV clades found in dogs showed no discernible correlation with prevalence or the temporal origins in the host. Notably, the Asian clade in dogs exhibited the highest nucleotide diversity, as indicated by the Pi (π) values (S3 Table). This particular clade, present in dog hosts, stands out as one of the oldest identified clades after the Indian-Sub clade [44]. The occurrence of similar clades in multiple hosts in a country highlights the events of cross-species transmission. However, currently, only phylogeny-based methods are available to identify clades; we believe the clade-defining SNPs reported in this study will serve as an alternative approach for clade identification, which will aid in the tracking of RABV clades.

Rabies is known to persist in two distinct forms: furious form and paralytic form. Hueffer et al. (2017) reported that the glycoprotein can modify host behaviour through a region that binds to the AchR receptor [45]. We find several mutations in an area of G protein (snake-toxin-like region, 175-203 residues in Glycoprotein) that binds to AchR. Most of the mutations in this region are universal (V181I, A182T, N177K). However, a stabilising mutation (V183I) is only present in Arctic clade sequences, further supporting the idea that clade-specific mutations may influence viral fitness and adaptation. The study of mutations also becomes important for understanding the differences in the fitness of the virus strains [46].

While this study encompassed sequences from 7 distinct hosts, dog hosts constituted more than half of the dataset. Countries significantly affected by RABV reported less WGS, impeding an in-depth understanding of RABV in these regions. The low-frequency mutations are not described, but these mutations are important for a comprehensive understanding of the virus evolution and circulation pattern. Furthermore, the inability to validate the functional implications of these pivotal mutations through wet lab experimentation remains a notable limitation. The diversity and distinct mutational repertoire of clades make evaluating vaccine efficacy against RABV clades crucial. Addressing these points would enhance our understanding of RABV evolution and facilitate more targeted and effective control measures against this global health concern.

## Acknowledgements

The authors would like the whole Utpal Tatu lab for their discussions.

## Author contributions

Conceptualisation: Ankeet Kumar and Utpal Tatu.

Data curation: Ankeet Kumar.

Formal analysis: Ankeet Kumar and Yashas Devasurmutt.

Investigation: Ankeet Kumar and Utpal Tatu.

Methodology: Ankeet Kumar and Utpal Tatu.

Supervision: Utpal Tatu.

Validation: Ankeet Kumar, Sheetal Tushir and Utpal Tatu.

Writing – original draft: Ankeet Kumar and Sujith S Nath.

Writing – review & editing: Ankeet Kumar, Sheetal Tushir and Utpal Tatu.

## Data Availability

The data supporting the findings in the study is available in the manuscript and as supporting data.

## Ethics approval

Not applicable

## Supplementary Data

**S1 File.** The list of genome accession utilised in the study with clade information.

**S2 File.** The modified script used to translate nucleotide variations (SNPs) to amino acids.

**S3 File.** A gff3 file with the information of coding regions for five proteins of RABV.

**S4 File.** An R script employed to merge the rows with nucleotide variation corresponding to the same amino acid.

**S5 File.** Host-wise characterisation of total mutations.

**S6 File.** A list of total and universal mutations represented in different clades of RABV.

**S7 File.** A list of clade-specific and clade-defining mutations for different hosts.

**S8 File.** Clade-specific mutations for the Cosmopolitan clade at the subclade level.

**S1 Table.** Count of mutations in different clades across hosts. S2 Table. Energy change values for mutations in the G protein. S3 Table. A summary table for clades found in Dogs.

**S4 Table.** A table representing diversity for the clades found in dogs.

## Supplementary Figures

**Fig S1.**
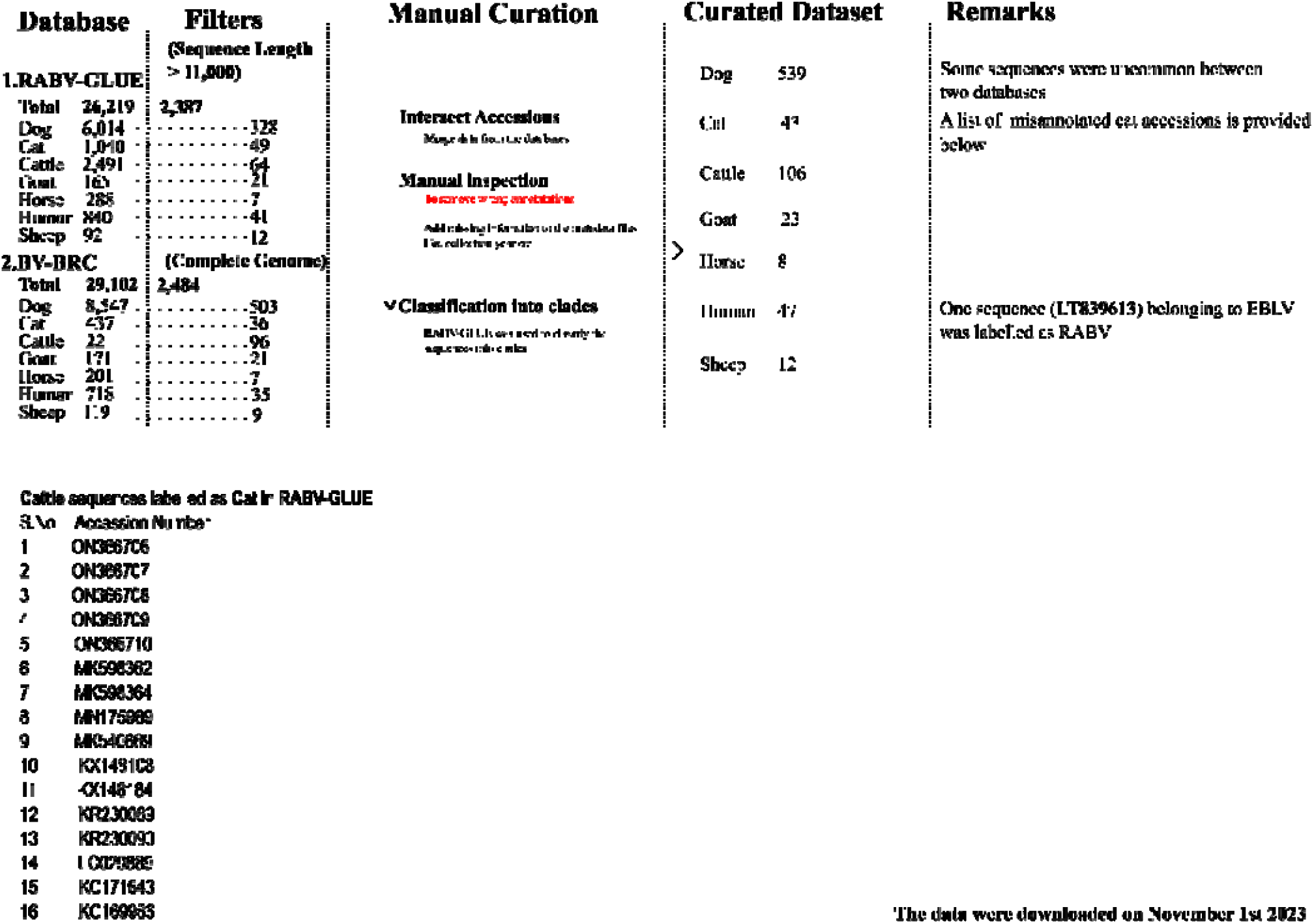
The pipeline used to curate the RABV dataset. The number of sequences is shown with the corresponding hosts and databases. The manual curation was done to identify the wrong annotations and add clade-related information.

**Fig S2.**
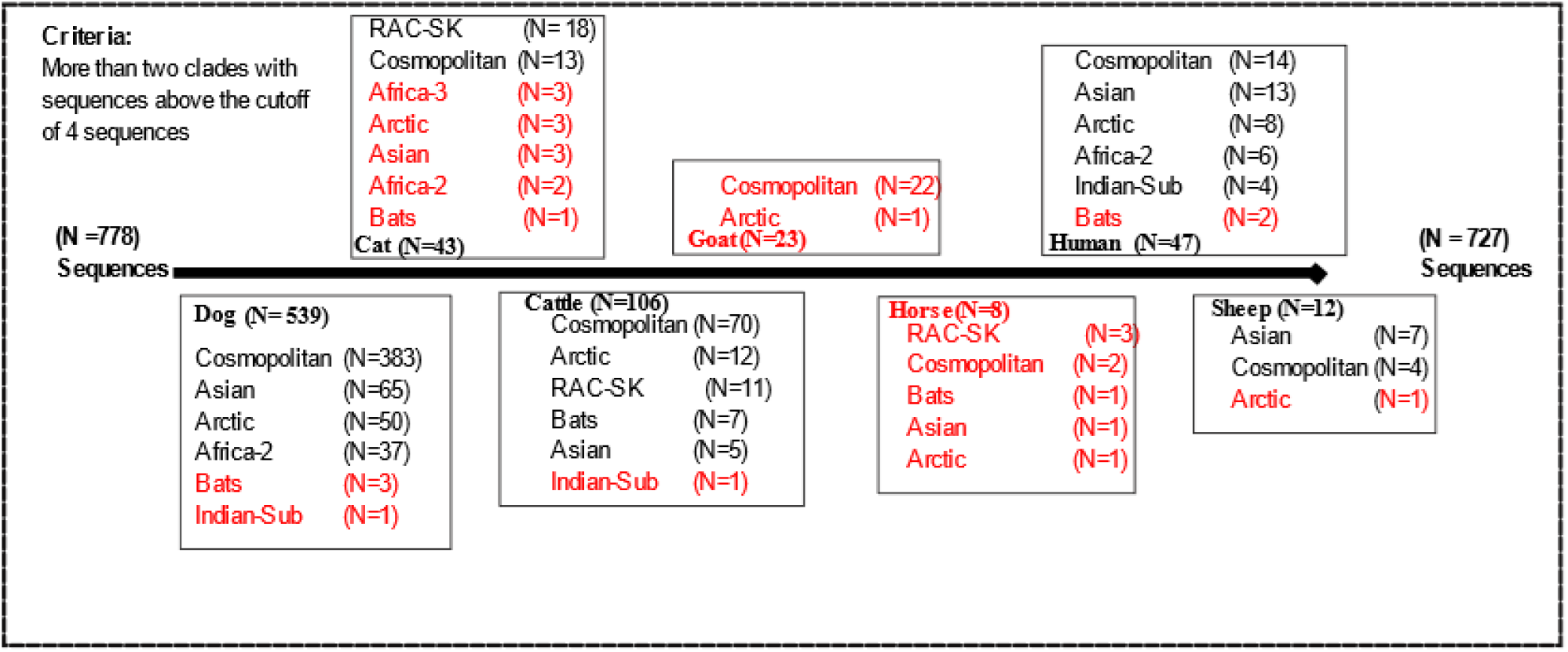
The hosts and clades used for high-frequency mutation analysis. The clades that were used for the analysis (> 4 sequences) are represented in black. The clades and hosts that are shown in red were not used for the analysis.

