## Supplementary material for "Identification of clade-defining single nucleotide polymorphisms for improved rabies virus surveillance": S1 Table mutation count.docx

**Cat Host**

| Major Clade | Annotation | Genome Count | SNP | SNP  silent | Deletion  frameshift | extragenic | Insertion  frameshift |
| --- | --- | --- | --- | --- | --- | --- | --- |
| Africa-2 | Glycoprotein | 2 | 87 | 331 | 0 | 0 | 0 |
| Africa-2 | Matrix Protein | 2 | 21 | 120 | 2 | 0 | 2 |
| Africa-2 | Nucleoprotein | 2 | 20 | 308 | 0 | 0 | 0 |
| Africa-2 | Phosphoprotein | 2 | 50 | 184 | 0 | 0 | 0 |
| Africa-2 | RdRp | 2 | 155 | 1458 | 7 | 0 | 6 |
| Africa-2 | Extragenic | 2 | 0 | 0 | 0 | 225 | 0 |
| Africa-3 | Glycoprotein | 3 | 150 | 425 | 4 | 0 | 4 |
| Africa-3 | Matrix Protein | 3 | 47 | 150 | 4 | 0 | 2 |
| Africa-3 | Nucleoprotein | 3 | 54 | 417 | 0 | 0 | 3 |
| Africa-3 | Phosphoprotein | 3 | 96 | 248 | 0 | 0 | 0 |
| Africa-3 | RdRp | 3 | 267 | 1951 | 14 | 0 | 13 |
| Africa-3 | Extragenic | 3 | 0 | 0 | 0 | 353 | 0 |
| Arctic | Glycoprotein | 3 | 134 | 425 | 0 | 0 | 0 |
| Arctic | Matrix Protein | 3 | 36 | 154 | 0 | 0 | 0 |
| Arctic | Nucleoprotein | 3 | 39 | 375 | 0 | 0 | 0 |
| Arctic | Phosphoprotein | 3 | 91 | 252 | 9 | 0 | 8 |
| Arctic | RdRp | 3 | 291 | 1957 | 9 | 0 | 9 |
| Arctic | Extragenic | 3 | 0 | 0 | 0 | 634 | 0 |
| Asian | Glycoprotein | 3 | 121 | 562 | 2 | 0 | 1 |
| Asian | Matrix Protein | 3 | 33 | 204 | 0 | 0 | 0 |
| Asian | Nucleoprotein | 3 | 32 | 471 | 0 | 0 | 1 |
| Asian | Phosphoprotein | 3 | 85 | 309 | 7 | 0 | 5 |
| Asian | RdRp | 3 | 268 | 2460 | 13 | 0 | 12 |
| Asian | Extragenic | 3 | 0 | 0 | 0 | 336 | 0 |
| Bats | Glycoprotein | 1 | 65 | 183 | 0 | 0 | 0 |
| Bats | Matrix Protein | 1 | 21 | 71 | 0 | 0 | 0 |
| Bats | Nucleoprotein | 1 | 19 | 159 | 0 | 0 | 0 |
| Bats | Phosphoprotein | 1 | 43 | 106 | 2 | 0 | 2 |
| Bats | RdRp | 1 | 115 | 845 | 9 | 0 | 8 |
| Bats | Extragenic | 1 | 0 | 0 | 0 | 116 | 0 |
| Cosmopolitan | Glycoprotein | 13 | 481 | 1212 | 1 | 0 | 1 |
| Cosmopolitan | Matrix Protein | 13 | 140 | 424 | 0 | 0 | 0 |
| Cosmopolitan | Nucleoprotein | 13 | 128 | 1137 | 0 | 0 | 2 |
| Cosmopolitan | Phosphoprotein | 13 | 276 | 773 | 1 | 0 | 1 |
| Cosmopolitan | RdRp | 13 | 863 | 5224 | 17 | 0 | 15 |
| Cosmopolitan | Extragenic | 13 | 0 | 0 | 0 | 1819 | 0 |
| RAC-SK | Nucleoprotein | 18 | 420 | 3375 | 2 | 0 | 1 |
| RAC-SK | Phosphoprotein | 18 | 221 | 667 | 16 | 0 | 8 |
| RAC-SK | Extragenic | 18 | 0 | 0 | 0 | 479 | 0 |

**Cattle Host**

| Major Clade | Annotation | Genome Count | SNP | SNP  silent | Deletion  frameshift | extragenic | Insertion  frameshift |
| --- | --- | --- | --- | --- | --- | --- | --- |
| Arctic | Glycoprotein | 12 | 526 | 1709 | 0 | 0 | 2 |
| Arctic | Matrix Protein | 12 | 135 | 666 | 0 | 0 | 0 |
| Arctic | Nucleoprotein | 12 | 148 | 1503 | 0 | 0 | 0 |
| Arctic | Phosphoprotein | 12 | 349 | 1051 | 18 | 0 | 18 |
| Arctic | RdRp | 12 | 1154 | 7866 | 34 | 0 | 34 |
| Arctic | Extragenic | 12 | 0 | 0 | 0 | 2391 | 0 |
| Asian | Glycoprotein | 4 | 188 | 724 | 16 | 0 | 12 |
| Asian | Matrix Protein | 4 | 49 | 237 | 4 | 0 | 4 |
| Asian | Nucleoprotein | 4 | 40 | 613 | 0 | 0 | 0 |
| Asian | Phosphoprotein | 4 | 134 | 382 | 4 | 0 | 4 |
| Asian | RdRp | 4 | 349 | 3232 | 24 | 0 | 20 |
| Asian | Extragenic | 4 | 0 | 0 | 0 | 380 | 0 |
| Bats | Glycoprotein | 7 | 157 | 665 | 6 | 0 | 6 |
| Bats | Matrix Protein | 7 | 88 | 306 | 0 | 0 | 0 |
| Bats | Nucleoprotein | 7 | 142 | 1236 | 4 | 0 | 2 |
| Bats | Phosphoprotein | 7 | 51 | 177 | 0 | 0 | 0 |
| Bats | RdRp | 7 | 123 | 881 | 13 | 0 | 12 |
| Bats | Extragenic | 7 | 0 | 0 | 0 | 409 | 0 |
| Cosmopolitan | Glycoprotein | 70 | 2437 | 6113 | 1 | 0 | 2 |
| Cosmopolitan | Matrix Protein | 70 | 739 | 2139 | 0 | 0 | 0 |
| Cosmopolitan | Nucleoprotein | 70 | 702 | 5719 | 1 | 0 | 6 |
| Cosmopolitan | Phosphoprotein | 70 | 1405 | 3994 | 13 | 0 | 10 |
| Cosmopolitan | RdRp | 70 | 4602 | 26976 | 82 | 0 | 80 |
| Cosmopolitan | Extragenic | 70 | 0 | 0 | 0 | 9713 | 0 |
| Indian-Sub | Glycoprotein | 1 | 42 | 186 | 0 | 0 | 0 |
| Indian-Sub | Matrix Protein | 1 | 12 | 64 | 0 | 0 | 0 |
| Indian-Sub | Nucleoprotein | 1 | 11 | 152 | 0 | 0 | 0 |
| Indian-Sub | Phosphoprotein | 1 | 30 | 105 | 2 | 0 | 1 |
| Indian-Sub | RdRp | 1 | 80 | 804 | 6 | 0 | 6 |
| Indian-Sub | Extragenic | 1 | 0 | 0 | 0 | 110 | 0 |
| RAC-SK | Nucleoprotein | 11 | 253 | 2059 | 0 | 0 | 0 |
| RAC-SK | Phosphoprotein | 11 | 191 | 465 | 17 | 0 | 8 |
| RAC-SK | RdRp | 11 | 124 | 968 | 4 | 0 | 4 |
| RAC-SK | Extragenic | 11 | 0 | 0 | 0 | 300 | 0 |

**Dog Host**

| Major Clade | Annotation | Genome Count | SNP | SNP  silent | Deletion  frameshift | extragenic | Insertion  frameshift |
| --- | --- | --- | --- | --- | --- | --- | --- |
| Africa-2 | Glycoprotein | 37 | 1608 | 6063 | 10 | 0 | 8 |
| Africa-2 | Matrix Protein | 37 | 393 | 2230 | 37 | 0 | 37 |
| Africa-2 | Nucleoprotein | 37 | 393 | 5536 | 6 | 0 | 7 |
| Africa-2 | Phosphoprotein | 37 | 941 | 3586 | 2 | 0 | 1 |
| Africa-2 | RdRp | 37 | 2789 | 26916 | 100 | 0 | 91 |
| Africa-2 | Extragenic | 37 | 0 | 0 | 0 | 3885 | 0 |
| Arctic | Glycoprotein | 50 | 2111 | 7168 | 4 | 0 | 16 |
| Arctic | Matrix Protein | 50 | 573 | 2620 | 0 | 0 | 1 |
| Arctic | Nucleoprotein | 50 | 649 | 6206 | 4 | 0 | 4 |
| Arctic | Phosphoprotein | 50 | 1403 | 4209 | 99 | 0 | 82 |
| Arctic | RdRp | 50 | 4331 | 31938 | 147 | 0 | 148 |
| Arctic | Extragenic | 50 | 0 | 0 | 0 | 9202 | 0 |
| Asian | Glycoprotein | 65 | 2834 | 12020 | 113 | 0 | 85 |
| Asian | Matrix Protein | 65 | 786 | 4237 | 30 | 0 | 29 |
| Asian | Nucleoprotein | 65 | 745 | 10037 | 1 | 0 | 10 |
| Asian | Phosphoprotein | 65 | 2081 | 6459 | 82 | 0 | 75 |
| Asian | RdRp | 65 | 5646 | 51413 | 365 | 0 | 323 |
| Asian | Extragenic | 65 | 0 | 0 | 0 | 6655 | 0 |
| Bats | Nucleoprotein | 3 | 61 | 529 | 0 | 0 | 0 |
| Bats | Phosphoprotein | 3 | 32 | 85 | 1 | 0 | 0 |
| Bats | Extragenic | 3 | 0 | 0 | 0 | 90 | 0 |
| Cosmopolitan | Glycoprotein | 383 | 12938 | 35531 | 9 | 0 | 13 |
| Cosmopolitan | Matrix Protein | 383 | 4005 | 12040 | 0 | 0 | 0 |
| Cosmopolitan | Nucleoprotein | 383 | 3743 | 32320 | 3 | 0 | 3 |
| Cosmopolitan | Phosphoprotein | 383 | 7782 | 22912 | 23 | 0 | 23 |
| Cosmopolitan | RdRp | 383 | 24877 | 153083 | 448 | 0 | 441 |
| Cosmopolitan | Extragenic | 383 | 0 | 0 | 0 | 54795 | 0 |
| Indian-Sub | Glycoprotein | 1 | 40 | 190 | 0 | 0 | 0 |
| Indian-Sub | Matrix Protein | 1 | 13 | 64 | 0 | 0 | 0 |
| Indian-Sub | Nucleoprotein | 1 | 12 | 148 | 0 | 0 | 0 |
| Indian-Sub | Phosphoprotein | 1 | 32 | 103 | 3 | 0 | 2 |
| Indian-Sub | RdRp | 1 | 81 | 815 | 5 | 0 | 5 |
| Indian-Sub | Extragenic | 1 | 0 | 0 | 0 | 107 | 0 |

**Goat Host**

| Major Clade | Annotation | Genome Count | SNP | SNP  silent | Deletion  frameshift | extragenic | Insertion  frameshift |
| --- | --- | --- | --- | --- | --- | --- | --- |
| Arctic | Glycoprotein | 1 | 41 | 146 | 0 | 0 | 1 |
| Arctic | Matrix Protein | 1 | 11 | 52 | 0 | 0 | 0 |
| Arctic | Nucleoprotein | 1 | 13 | 129 | 0 | 0 | 0 |
| Arctic | Phosphoprotein | 1 | 24 | 81 | 0 | 0 | 0 |
| Arctic | RdRp | 1 | 79 | 634 | 4 | 0 | 4 |
| Arctic | Extragenic | 1 | 0 | 0 | 0 | 189 | 0 |
| Cosmopolitan | Glycoprotein | 22 | 785 | 2036 | 0 | 0 | 0 |
| Cosmopolitan | Matrix Protein | 22 | 240 | 682 | 0 | 0 | 0 |
| Cosmopolitan | Nucleoprotein | 22 | 233 | 1885 | 0 | 0 | 0 |
| Cosmopolitan | Phosphoprotein | 22 | 466 | 1274 | 1 | 0 | 1 |
| Cosmopolitan | RdRp | 22 | 1489 | 8657 | 22 | 0 | 22 |
| Cosmopolitan | Extragenic | 22 | 0 | 0 | 0 | 3149 | 0 |

**Horse Host**

| Major Clade | Annotation | Genome Count | SNP | SNP  silent | Deletion  frameshift | extragenic | Insertion  frameshift |
| --- | --- | --- | --- | --- | --- | --- | --- |
| Arctic | Glycoprotein | 1 | 41 | 140 | 0 | 0 | 1 |
| Arctic | Matrix Protein | 1 | 11 | 55 | 0 | 0 | 0 |
| Arctic | Nucleoprotein | 1 | 12 | 129 | 0 | 0 | 0 |
| Arctic | Phosphoprotein | 1 | 24 | 87 | 0 | 0 | 0 |
| Arctic | RdRp | 1 | 69 | 642 | 3 | 0 | 3 |
| Arctic | Extragenic | 1 | 0 | 0 | 0 | 175 | 0 |
| Asian | Glycoprotein | 1 | 46 | 181 | 4 | 0 | 3 |
| Asian | Matrix Protein | 1 | 13 | 59 | 1 | 0 | 1 |
| Asian | Nucleoprotein | 1 | 10 | 155 | 0 | 0 | 0 |
| Asian | Phosphoprotein | 1 | 32 | 98 | 1 | 0 | 1 |
| Asian | RdRp | 1 | 86 | 807 | 6 | 0 | 5 |
| Asian | Extragenic | 1 | 0 | 0 | 0 | 96 | 0 |
| Bats | Glycoprotein | 1 | 63 | 185 | 1 | 0 | 1 |
| Bats | Matrix Protein | 1 | 22 | 73 | 0 | 0 | 0 |
| Bats | Nucleoprotein | 1 | 19 | 156 | 0 | 0 | 0 |
| Bats | Phosphoprotein | 1 | 43 | 105 | 2 | 0 | 2 |
| Bats | RdRp | 1 | 112 | 851 | 7 | 0 | 6 |
| Bats | Extragenic | 1 | 0 | 0 | 0 | 124 | 0 |
| Cosmopolitan | Glycoprotein | 2 | 67 | 211 | 1 | 0 | 1 |
| Cosmopolitan | Matrix Protein | 2 | 24 | 68 | 0 | 0 | 0 |
| Cosmopolitan | Nucleoprotein | 2 | 21 | 196 | 0 | 0 | 0 |
| Cosmopolitan | Phosphoprotein | 2 | 51 | 124 | 0 | 0 | 0 |
| Cosmopolitan | RdRp | 2 | 140 | 883 | 2 | 0 | 2 |
| Cosmopolitan | Extragenic | 2 | 0 | 0 | 0 | 297 | 0 |
| RAC-SK | Nucleoprotein | 3 | 69 | 559 | 0 | 0 | 0 |
| RAC-SK | Phosphoprotein | 3 | 47 | 127 | 4 | 0 | 2 |
| RAC-SK | Extragenic | 3 | 0 | 0 | 0 | 78 | 0 |

**Humans**

| Major Clade | Annotation | Genome  Count | SNP | SNP  silent | Deletion  frameshift | extragenic | Insertion  frameshift |
| --- | --- | --- | --- | --- | --- | --- | --- |
| Africa-2 | Glycoprotein | 6 | 250 | 987 | 0 | 0 | 0 |
| Africa-2 | Matrix Protein | 6 | 64 | 368 | 6 | 0 | 6 |
| Africa-2 | Nucleoprotein | 6 | 64 | 896 | 1 | 0 | 1 |
| Africa-2 | Phosphoprotein | 6 | 151 | 583 | 0 | 0 | 0 |
| Africa-2 | RdRp | 6 | 471 | 4347 | 13 | 0 | 13 |
| Africa-2 | Extragenic | 6 | 0 | 0 | 0 | 673 | 0 |
| Arctic | Glycoprotein | 8 | 328 | 1130 | 0 | 0 | 5 |
| Arctic | Matrix Protein | 8 | 87 | 435 | 0 | 0 | 0 |
| Arctic | Nucleoprotein | 8 | 103 | 1035 | 0 | 0 | 0 |
| Arctic | Phosphoprotein | 8 | 221 | 689 | 2 | 0 | 2 |
| Arctic | RdRp | 8 | 613 | 5125 | 20 | 0 | 20 |
| Arctic | Extragenic | 8 | 0 | 0 | 0 | 1348 | 0 |
| Asian | Glycoprotein | 13 | 533 | 2362 | 19 | 0 | 12 |
| Asian | Matrix Protein | 13 | 160 | 875 | 7 | 0 | 5 |
| Asian | Nucleoprotein | 13 | 145 | 2004 | 0 | 0 | 3 |
| Asian | Phosphoprotein | 13 | 417 | 1298 | 11 | 0 | 11 |
| Asian | RdRp | 13 | 867 | 8057 | 48 | 0 | 44 |
| Asian | Extragenic | 13 | 0 | 0 | 0 | 1240 | 0 |
| Bats | Glycoprotein | 2 | 88 | 337 | 6 | 0 | 5 |
| Bats | Matrix Protein | 2 | 25 | 71 | 0 | 0 | 0 |
| Bats | Nucleoprotein | 2 | 43 | 349 | 1 | 0 | 2 |
| Bats | Phosphoprotein | 2 | 69 | 139 | 6 | 0 | 5 |
| Bats | RdRp | 2 | 115 | 873 | 12 | 0 | 11 |
| Bats | Extragenic | 2 | 0 | 0 | 0 | 176 | 0 |
| Cosmopolitan | Glycoprotein | 14 | 479 | 1365 | 1 | 0 | 1 |
| Cosmopolitan | Matrix Protein | 14 | 145 | 441 | 0 | 0 | 0 |
| Cosmopolitan | Nucleoprotein | 14 | 145 | 1190 | 2 | 0 | 0 |
| Cosmopolitan | Phosphoprotein | 14 | 278 | 824 | 2 | 0 | 2 |
| Cosmopolitan | RdRp | 14 | 940 | 5904 | 24 | 0 | 24 |
| Cosmopolitan | Extragenic | 14 | 0 | 0 | 0 | 2060 | 0 |
| Indian-Sub | Glycoprotein | 4 | 130 | 729 | 2 | 0 | 2 |
| Indian-Sub | Matrix Protein | 4 | 51 | 271 | 0 | 0 | 0 |
| Indian-Sub | Nucleoprotein | 4 | 46 | 603 | 0 | 0 | 0 |
| Indian-Sub | Phosphoprotein | 4 | 128 | 420 | 8 | 0 | 5 |
| Indian-Sub | RdRp | 4 | 318 | 3240 | 24 | 0 | 24 |
| Indian-Sub | Extragenic | 4 | 0 | 0 | 0 | 408 | 0 |

**Sheep Host**

| Major Clade | Annotation | Genome count | SNP | SNP  silent | Deletion  frameshift | extragenic | Insertion frameshift |
| --- | --- | --- | --- | --- | --- | --- | --- |
| Arctic | Glycoprotein | 4 | 178 | 559 | 0 | 0 | 0 |
| Arctic | Matrix Protein | 4 | 50 | 194 | 0 | 0 | 0 |
| Arctic | Nucleoprotein | 4 | 57 | 498 | 0 | 0 | 0 |
| Arctic | Phosphoprotein | 4 | 124 | 337 | 12 | 0 | 12 |
| Arctic | RdRp | 4 | 381 | 2599 | 12 | 0 | 12 |
| Arctic | Extragenic | 4 | 0 | 0 | 0 | 838 | 0 |
| Asian | Glycoprotein | 1 | 48 | 181 | 4 | 0 | 3 |
| Asian | Matrix Protein | 1 | 12 | 58 | 1 | 0 | 1 |
| Asian | Nucleoprotein | 1 | 10 | 155 | 0 | 0 | 0 |
| Asian | Phosphoprotein | 1 | 33 | 96 | 1 | 0 | 1 |
| Asian | RdRp | 1 | 86 | 813 | 6 | 0 | 5 |
| Asian | Extragenic | 1 | 0 | 0 | 0 | 96 | 0 |
| Cosmopolitan | Glycoprotein | 7 | 250 | 623 | 0 | 0 | 0 |
| Cosmopolitan | Matrix Protein | 7 | 70 | 230 | 0 | 0 | 0 |
| Cosmopolitan | Nucleoprotein | 7 | 71 | 622 | 0 | 0 | 0 |
| Cosmopolitan | Phosphoprotein | 7 | 140 | 408 | 4 | 0 | 3 |
| Cosmopolitan | RdRp | 7 | 470 | 2777 | 13 | 0 | 11 |
| Cosmopolitan | Extragenic | 7 | 0 | 0 | 0 | 1006 | 0 |
