## Supplementary material for "Identification of clade-defining single nucleotide polymorphisms for improved rabies virus surveillance": S2 File_R script.docx

```
## By Federico M. Giorgi – Orignal Author
## Modified by – Ankeet Kumar

###### BASH section
### Download RABV sequences in FASTA format
input=input.fasta
### Run nucmer to obtain variant file
ref=NC001542.fasta # NCBI Reference Rabies
dos2unix $input
nucmer --forward -p nucmer $ref $input
show-coords -r -c -l nucmer.delta > nucmer.coords
show-snps nucmer.delta -T -l > nucmer.snps

###### R section
nucmer <- read.delim("nucmer.snps", as.is = TRUE, skip = 4, header = FALSE, sep = "\t")
colnames(nucmer) <- c("refpos", "refvar", "qvar", "qpos", "", "", "", "", "rlength", "qlength", "", "", "rname", "qname")
rownames(nucmer) <- paste0("var", 1:nrow(nucmer))

# Fix IUPAC codes
table(nucmer$qvar)
nucmer<-nucmer[!nucmer$qvar%in%c("B","D","H","K","M","N","R","S","V","W","Y"),]
nrow(nucmer)

### Aminoacid variant list ----
# Load reference sequence
library(seqinr)
library(Biostrings)
refseq<-read.fasta("NC001542_refseq.fasta",forceDNAtolower=FALSE)[[1]]

# Load GFF3
gff3<-read.delim("NC_001542_gff3.gff3",as.is=TRUE,skip=2,header=FALSE)
annot<-setNames(gff3[,10],gff3[,9])

header<-c("sample","refpos","refvar","qvar","qpos","qlength","protein","variant","varclass","annotation")
results<-matrix(NA,ncol=length(header),nrow=0)
colnames(results)<-header
samples<-unique(nucmer$qname)
pb<-txtProgressBar(0,length(samples),style=3)
for (pbi in 1:length(samples)){ # This will update the nucmer object
  sample<-samples[pbi]
  allvars<-nucmer[nucmer$qname==sample,]
  # Check changes in query protein sequence according to variants
  for(i in 1:nrow(allvars)){ # Assuming they are sorted numerically
    nucline<-allvars[i,]
    refpos<-nucline[1,"refpos"]
    refvar<-nucline[1,"refvar"]
    qvar<-nucline[1,"qvar"]
    qpos<-nucline[1,"qpos"]
    qlength<-nucline[1,"qlength"]

    # Match over GFF3 annotation
    a<-refpos-gff3[,4]
    b<-refpos-gff3[,5]
    signs<-sign(a)*sign(b)
    w<-which(signs==-1)


    # Outside genes scenarios
    if(length(w)==0){
      if(refpos<gff3[1,4]){
        protein<-"5'UTR";output<-c(refpos,"extragenic")
      } else if(refpos>gff3[1,5]){
        protein<-"3'UTR";output<-c(refpos,"extragenic")
      } else {
        protein<-"intergenic";output<-c(refpos,"extragenic")
      }

    } else{ # Inside genes scenario
      start<-gff3[w,4]
      end<-gff3[w,5]
      protein<-gff3[w,9]
      refdnaseq<-Biostrings::DNAString(paste0(refseq[start:end],collapse=""))
      refpepseq<-Biostrings::translate(refdnaseq)
      refpepseq<-strsplit(as.character(refpepseq),"")[[1]]
      if(qvar=="."){ # Deletion scenario
        if((nchar(refvar)%%3)!=0){ # Deletion frameshift scenario
          mutpos<-ceiling((refpos-start+1)/3)
          output<-c(paste0(refpepseq[mutpos],mutpos),"deletion_frameshift")
        } else { # In-frame deletion
          varseq<-refseq
          varseq<-varseq[-(refpos:(refpos+nchar(refvar)-1))]
          varseq<-varseq[start:(end-nchar(refvar))]
          vardnaseq<-Biostrings::DNAString(paste0(varseq,collapse=""))
          varpepseq<-Biostrings::translate(vardnaseq)
          varpepseq<-strsplit(as.character(varpepseq),"")[[1]]

          for(j in 1:length(refpepseq)){
            refj<-refpepseq[j]
            varj<-varpepseq[j]
            if(refj!=varj){
              if(varj=="*"){
                output<-c(paste0(refj,j),"deletion_stop")
              } else {
                output<-c(paste0(refj,j),"deletion")
              }
              break()
            }
          }
        }
      } else if(refvar=="."){ # Insertion scenario
        if((nchar(qvar)%%3)!=0){ # Insertion frameshift scenario
          mutpos<-ceiling((refpos-start+1)/3)
          output<-c(paste0(refpepseq[mutpos],mutpos),"insertion_frameshift")
        } else { # In-frame insertion
          varseq<-c(refseq[1:refpos],strsplit(qvar,"")[[1]],refseq[(refpos+1):length(refseq)])
          varseq<-varseq[start:(end+nchar(qvar))]
          vardnaseq<-Biostrings::DNAString(paste0(varseq,collapse=""))
          varpepseq<-Biostrings::translate(vardnaseq)
          varpepseq<-strsplit(as.character(varpepseq),"")[[1]]

          for(j in 1:length(refpepseq)){
            refj<-refpepseq[j]
            varj<-varpepseq[j]
            if(refj!=varj){
              nr_aa_inserted<-nchar(qvar)/3
              multivarj<-varpepseq[j:(j+nr_aa_inserted-1)]
              if(any(multivarj=="*")){
                multivarj<-paste0(multivarj,collapse="")
                output<-c(paste0(multivarj,j),"insertion_stop")
              } else{
                multivarj<-paste0(multivarj,collapse="")
                output<-c(paste0(multivarj,j),"insertion")
              }
              break()
            }
          }
        }
      } else {
        if (nchar(qvar) == 1) {
          codonpos <- ceiling(((refpos+1) - start) / 3)
          codonstart <- (3*codonpos +(start)-3)
          codonend <- codonstart + 2
          allvars_codon <- allvars[allvars$refpos >= codonstart & allvars$refpos <= codonend, ]
          # If there are gaps in the translated sequence
          if (any(allvars_codon$qvar == ".")) {
            allvar_pos <- allvars_codon[(allvars_codon$qvar != "."),]
            allvar_sites <- allvar_pos$refpos
            after_changes <-allvar_pos$qvar
            varseq[allvar_sites] <- paste(after_changes)
            vardnaseq <- Biostrings::DNAString(paste0(varseq[(start):end], collapse=""))
            varpepseq <- Biostrings::translate(vardnaseq)
            varpepseq <- strsplit(as.character(varpepseq), "")[[1]]
            refdnaseq<-Biostrings::DNAString(paste0(refseq[(start):end],collapse=""))
            refpepseq<-Biostrings::translate(refdnaseq)
            refpepseq<-strsplit(as.character(refpepseq),"")[[1]]
            refaa <- refpepseq[codonpos]
            varaa <- varpepseq[codonpos]
            output <- c(paste0(refaa, codonpos, varaa), "deletion_frameshift")
          }else {
            if (any(allvars_codon$qpos != ".")){
              allvar_sites <- allvars_codon$refpos
              after_changes <- allvars_codon$qvar
              varseq <- refseq
              varseq[allvar_sites] <- paste(after_changes)
              vardnaseq <- Biostrings::DNAString(paste0(varseq[(start):end], collapse=""))
              varpepseq <- Biostrings::translate(vardnaseq)
              varpepseq <- strsplit(as.character(varpepseq), "")[[1]]
              refdnaseq<-Biostrings::DNAString(paste0(refseq[start:end],collapse=""))
              refpepseq<-Biostrings::translate(refdnaseq)
              refpepseq<-strsplit(as.character(refpepseq),"")[[1]]
              refaa <- refpepseq[codonpos]
              varaa <- varpepseq[codonpos]
              if (refaa[1]==varaa[1]) {
                refaa <- refpepseq[codonpos]
                varaa <- varpepseq[codonpos]
                output <- c(paste0(refaa, codonpos, varaa), "SNP_silent")
                # If the amino acid changes, output "SNP" or "SNP_stop"
              } else {
                if (varaa == "*") {
                  output <- c(paste0(refaa, codonpos, varaa), "SNP_stop")
                } else {
                  output <- c(paste0(refaa, codonpos, varaa), "SNP")
                }
              }
            }
          }
        }
      }
    }
    results <- rbind(results,c(sample,refpos,refvar,qvar,qpos,qlength,protein,output,annot[protein]))
  }
  setTxtProgressBar(pb,pbi)
}
write.csv(results, "results.csv", row.names = FALSE)
```
