## Supplementary material for "Identification of clade-defining single nucleotide polymorphisms for improved rabies virus surveillance": S2 Table G protein stabilisation values.docx

| **Mutation** | **Energy** | **Remark** | **Class** |
| --- | --- | --- | --- |
| **V183I** | 0.495 | Stabilising | Clade Specific Arctic |
| **L16M** | 0.225 | Stabilising | Clade Specific Arctic |
| **N355D** | 0.052 | Stabilising | Clade Specific Africa-2 |
| **F8V** | 0.405 | Stabilising | Clade Specific Africa-2 |
| **N375H** | 0.09 | Stabilising | Clade Specific Africa-2 |
| **Q263R** | -0.289 | Stabilising | Clade Specific Africa-2 |
| **N484K** | -0.174 | De-Stabilising | Clade Specific Africa-2 |
| E444K | 0.614 | Stabilising | Universal |
| V181I | 1.052 | Stabilising | Universal |
| N177K | 0.545 | Stabilising | Universal |
| I509V | 0.018 | Stabilising | Universal |
| E224K | 0.972 | Stabilising | Universal |
| R445Q | -0.205 | De-Stabilising | Universal |
| M206T | -0.273 | De-Stabilising | Universal |
| M75V | -0.636 | De-Stabilising | Universal |
| E499K | 0.007 | Stabilising | Universal |
| A182T | 1.046 | Stabilising | Universal |
| V408E | -0.078 | De-Stabilising | Universal |
| W480C | -0.567 | De-Stabilising | Universal |
| N389H | 0.523 | Stabilising | Universal |
| G274D | 0.141 | Stabilising | Universal |
| P504S | 0.806 | Stabilising | Universal |
| N266D | -0.154 | De-Stabilising | Universal |

**S1 Table. Different mutations observed in the G protein for Canine RABV.**
