## Supplementary material for "Identification of clade-defining single nucleotide polymorphisms for improved rabies virus surveillance": S3 File gff3 gene coordinates.docx

##sequence-region NC_001542.1 1 11932

##species https://www.ncbi.nlm.nih.gov/Taxonomy/Browser/wwwtax.cgi?id=11292

NC_001542.1	RefSeq	CDS	71	1423	Kumar A	+	.	N	Nucleoprotein

NC_001542.1	RefSeq	CDS	1514	2407	Kumar A	+	.	P	Phosphoprotein

NC_001542.1	RefSeq	CDS	2496	3104	Kumar A	+	.	M	Matrix Protein

NC_001542.1	RefSeq	CDS	3318	4892	Kumar A	+	.	G	Glycoprotein

NC_001542.1	RefSeq	CDS	5418	11846	Kumar A	+	.	L	RdRp
