## Supplementary material for "Identification of clade-defining single nucleotide polymorphisms for improved rabies virus surveillance": S3 Table Dog clade summary table.docx

| Major Clade | Total Mutation | SNP | SNP silent | Deletion Frameshift | Extragenic | Insertion Frameshift | Clade Specific mutation | Clade defining |
| --- | --- | --- | --- | --- | --- | --- | --- | --- |
| Africa-2 (N=37) | 54639 | 6124 | 44331 | 155 | 3885 | 144 | N_F213Y P_K139K, P_P159L M_S185S G_F8V, G_Q263R,G_N355D, G_N375H, G_N484K L_M93I, L_T436A, L_N683D, L_Q745Q, L_K1624R, L_T1961A | G_N375H, G_N484K, G_Q263R, L_E52D L_M93I,L_N683D,L_T1961A,L_T436A, N_F213Y,P_P159 |
| Arctic (N=50) | 70915 | 9067 | 52141 | 254 | 9202 | 251 | N_M60L, N_P135P P_E165D, P_A267G G_L16M, G_V183I | G_L16M,N_M60L,P_A267G |
| Asian (N=65) | 104026 | 12092 | 84166 | 591 | 6655 | 522 | L_L2I, L_L424I, L_A1535S, L_N1839E | L_L424I |
| Bats (N=1) | 798 | 93 | 614 | 1 | 90 | 0 | < Cutoff of 4 sequences | |
| Cosmopolitan (N=383) | 364989 | 53345 | 255886 | 483 | 54795 | 480 | No Clade specific mutation | |
| Indian-Sub (N=3) | 1620 | 178 | 1320 | 8 | 107 | 7 | < Cutoff of 4 sequences | |
