## Supplementary material for "Identification of clade-defining single nucleotide polymorphisms for improved rabies virus surveillance": S4 File merge results script.docx

\# Load the library required for merging script

library(dplyr)

\# load the metadata for particular host under analysis

metadata <- read.csv("metadata.csv")

\# Check accessions

unique(metadata$sample)

\# load the metadata for particular host under analysis

results<- read.csv("results.csv")

\# Check accessions

unique(results$sample)

setdiff(unique(metadata$sample), unique(results$sample))

\## If the value of setdiff is zero, then proceed, otherwise check your files

\# make a column called protvar, which shows protein and variant in a single column

\# this is to make sure that same variant present in other protein should not create

\# any error

results$protvar<-paste0(results$protein, "_",results$variant)

\# merge the results and metadata to contain inforamtion of clades with the sequences

results_clade <- merge(results, metadata, by= "sample")

\# save the result as csv file

write.csv(results_clade, "RESULTS_CLADE.csv", row.names = F)

\# convert the the numeric values to characteric, as the rows are going to merge

results_clade<- results_clade%>%

mutate(qpos = as.character(qpos),

qlength = as.character(qlength),

refpos = as.character(refpos))

\# produce a dataframe of merged results

merged_results_clade<- results_clade%>%

group_by(sample, variant, protein) %>%

filter(n() <= 3) %>%

reframe(

refpos = paste(refpos, collapse = "_"),

refvar = paste(refvar, collapse = ""),

qvar = paste(qvar, collapse = ""),

qpos = paste(qpos, collapse = "_"),

qlength = first(qlength),

varclass = first(varclass),

annotation = first(annotation),

majorclade = first(majorclade),

minorclade = first(minorclade),

country = first(country),

collectionyear = first(collectionyear),

host = first(host))

write.csv(merged_results_clade,"MERGED_RESULTS.csv", row.names = F)
