## Supplementary material for "Identification of clade-defining single nucleotide polymorphisms for improved rabies virus surveillance": S4 Table diversity in dog clades.docx

| **Clade** | **Number**  **Of sequences** | **S**^a^ | **h**^b^ | **Hd**^c^ | **K**^d^ | **Pi**^e^ | **PiJC**^f^ |
| --- | --- | --- | --- | --- | --- | --- | --- |
| Africa-2 | 37 | 1973 | 37 | 1.00000 | 299.87 | 0.02817 | 0.02889 |
| Arctic | 50 | 2681 | 49 | 0.99918 | 607.39 | 0.05706 | 0.06026 |
| Asian | 65 | 3636 | 65 | 1.00000 | 946.21 | 0.08889 | 0.09631 |
| Cosmopolitan | 379 | 4828 | 345 | 0.99929 | 610.30 | 0.05733 | 0.06009 |
| Bats | 3 | 266 | 3 | 1.00000 | 177.66 | 0.01669 | 0.01697 |

**S1 Table. The data nucleotide-based genetic diversity data was obtained using the DnaSP program**

^a^S, Number of segregating sites

^b^h, Number of haplotypes

^c^Hd, Haplotype diversity

^d^K, Average number of differences

^e^Pi, Nucleotide diversity

^f^PiJC, Nucleotide diversity with JC
