## Supplementary figures and images for "Identification of clade-defining single nucleotide polymorphisms for improved rabies virus surveillance"

### S1 Fig.tiff

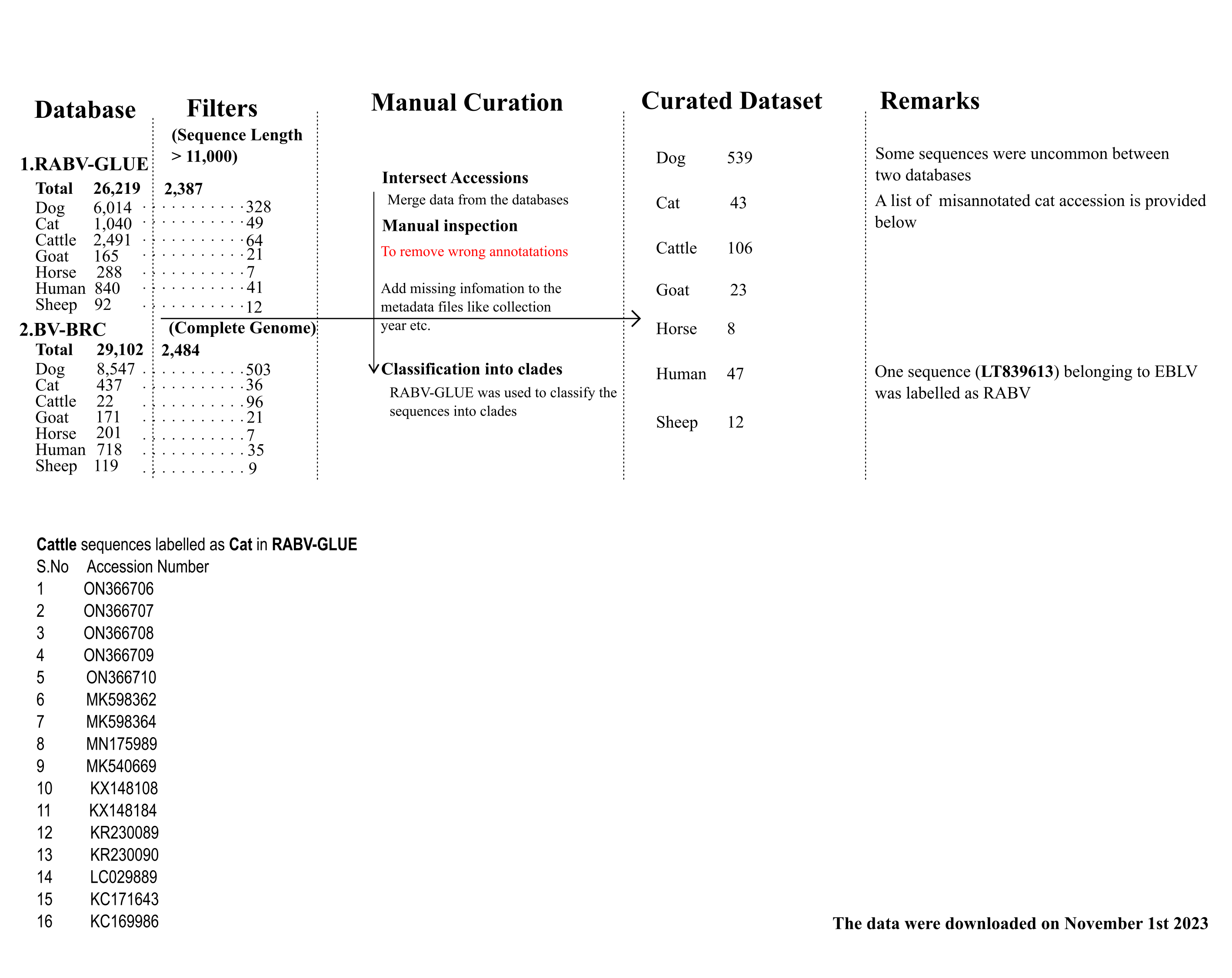

### S2 Fig.tiff

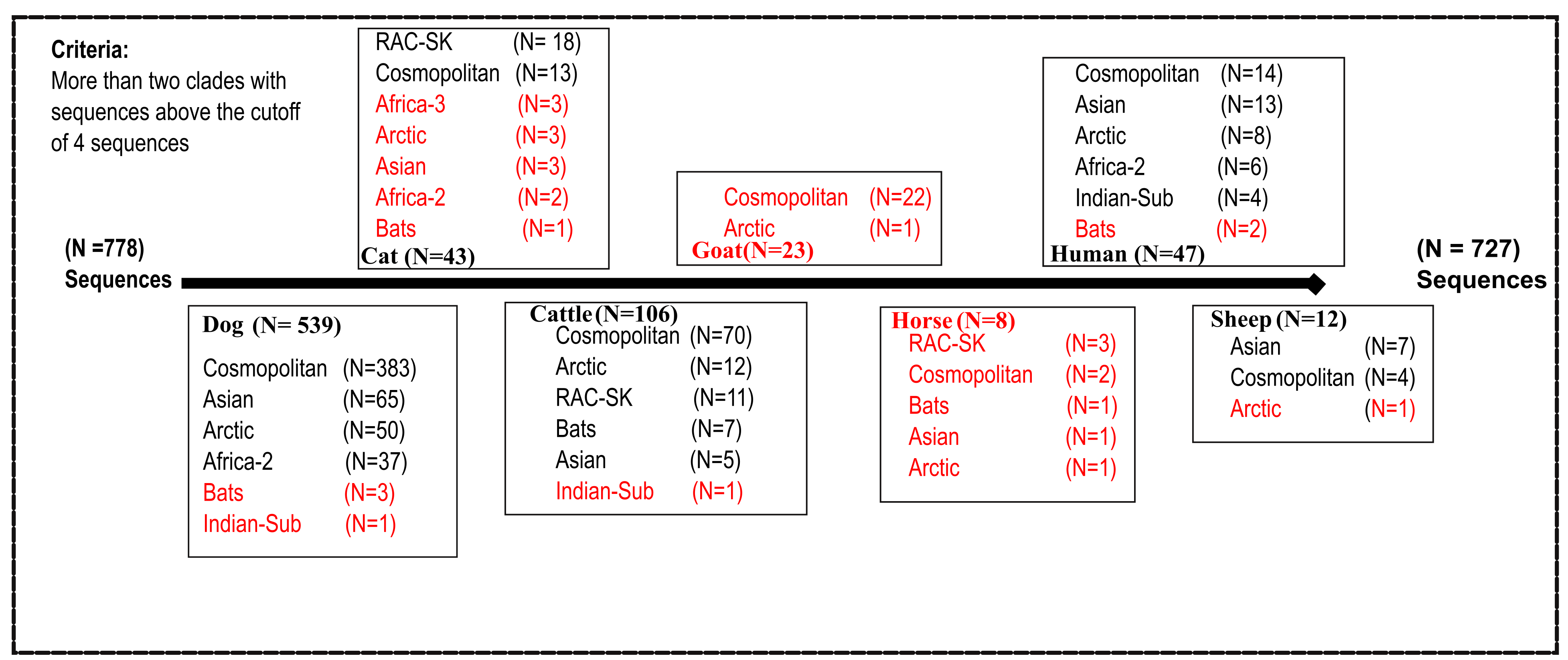
